# Enhancing Agrobacterium-Mediated Soybean Transformation Efficiency with an Auxiliary Solution

**DOI:** 10.1101/2024.04.19.590356

**Authors:** Luying Chen, Liang Wang, Yongguang Li, Shaojie Han

## Abstract

Soybean provides oil, protein, and biofuel. Efficient transformation systems are vital for advancing soybean research. Currently, *Agrobacterium rhizogenes-*mediated transformation is the predominant method for facilitating rapid transformation, producing transgenic hairy roots. However, the limitations of soybean transformation technology primarily originate from its low efficiency and genotype dependency, leaving significant room for improvement in the development more universally applicable and efficient methods. In this study, we explored the enhancement of soybean transformation efficiency through the generation and validation of three reporter vectors (ZsGreen, TdTomato, and Ruby) and the strategic use of *Agrobacterium* Auxiliary Solution (AAS) containing Silwet L-77 and hormone mixtures. Our findings demonstate that the incorporation of hormone mixtures and Silwet L-77 into AAS significantly improves hairy root transformation rates. Specifically, the combination of hormone mixtures with Silwet L-77 substantially increased both total root and cotyledon transformation efficiencies compared to the control. We also assessed the impact of vector size on transformation efficiency, observing a notable decrease in efficiency with larger vectors such as the Ruby cassette compared to smaller markers like GFP and RFP. Furthermore, our study examined the effects of AAS on the co-transformation rate of two separate vectors, revealing a slight but significant reduction in efficiency compared to single vector transformations. Additionally, we evaluated the role of AAS in enhancing soybean hypocotyl transformation rates in composite soybean plants across various varieties. The results consistently showed an increase in both positive roots and explant efficiencies with the addition of AAS, indicating its broad applicability and effectiveness in soybean transformation. However, significant varietal differences in transformation rates were observed, particularly between “Forrest” and other varieties such as “Williams 82” and “Dongnong 50”. In summary, our research emphasizes the significant role of auxiliary agents and vector size in optimizing soybean transformation techniques, providing valuable insights for future advancements in soybean genetic modification and biotechnological research.

## Introduction

Soybean (*Glycine max* (L.) Merr.), a globally significant economic crop, plays a critical role in providing essential oil and protein for human consumption and is also a key resource for biofuel production(Bai et al. 2020). The evolution of biotechnology has propelled the importance of advanced breeding techniques, functional studies, and precise genetic modifications in soybean research(Jin et al. 2021). Nonetheless, the narrow range of genetic diversity available for certain desired traits and the extended duration required for traditional breeding present considerable challenges in the conventional cultivation of soybeans(Lyzenga et al. 2021; Torkamaneh et al. 2021).

Soybean transformation, surpassing traditional breeding methods, plays an indispensable role in addressing fundamental biological queries(Yamada et al. 2012; Jin et al. 2021; Huang et al. 2022a). Despite the development and refinement of soybean transformation systems over the past three decades since their initial introduction(Yun et al. 2022; Zhai et al. 2022; Li et al. 2023), achieving stable genetic transformation in soybean remains less efficient compared to other crops(Cheng et al. 2021). This inefficiency is especially notable given the expanding genomic resources for soybeans and the urgency to effectively harness genome editing technologies for both biotechnological applications and fundamental research in soybeans. Consequently, there is a pressing need for more efficient transformation systems to propel soybean research forward. Presently, *Agrobacterium rhizogenes* (*A. rhizogenes*)-mediated transformation and biolistic methods stand as the predominant techniques employed in soybean transformation(Li et al. 2017a).

Utilizing *A. rhizogenes* facilitates the swift and straightforward transformation of soybeans, leading to the production of transgenic hairy roots. *A. rhizogenes*, a Gram-negative soil bacterium belonging to the *Agrobacterium* genus within the Rhizobiaceae family(Otten 2021), operates similarly to *Agrobacterium tumefaciens* (*A. tumefaciens*). It transfers its intrinsic T-DNA from root-inducing plasmids, extrachromosomal replicons, directly into the plant’s genomic DNA. The root-inducing pRi2659 A. tumefaciens Ri plasmid, present in A. tumefaciens, harbors root locus (rol) genes *rolA*, *rolB*, *rolC*, and *rolD* within the T-DNA region(Capone et al. 1989; Otten 2021). This plasmid can initiate the growth of hairy roots at damaged plant surfaces upon infection(Hanisch et al. 1990). This technique enables the quick generation of a substantial number of transgenic soybean roots in a relatively short timeframe, making them suitable for a variety of molecular assays and biological experiments(Huang et al. 2022b). These soybean hairy roots can undergo further differentiation, leading to the formation of new healing tissues and ultimately resulting in a heritable transformed line. Employing this method for overexpression or RNA interference (RNAi) of target genes is considerably less time-consuming compared to the genetic transformation of soybeans(Li et al. 2017b; Fan et al. 2020).

Despite their utility, these transformation techniques, particularly those involving *A. rhizogenes*, are often too inefficient and labor-intensive to fully satisfy the escalating demands of contemporary research. Presently, two primary *A. rhizogenes*-mediated soybean transformation methods are in use: Aerial root transformation(Matthews and Youssef 2016), which targets the soybean hypocotyl for transformation. This approach, however, results in a low ratio of positive (transgenic) roots relative to the total number of roots and is susceptible to a high incidence of false positives(Matthews and Youssef 2016; Fan et al. 2020). The second method is Aseptic histoculture(Chen et al. 2018; Cheng et al. 2021), where transformation occurs at the soybean cotyledon. This technique, while innovative, faces challenges such as being cumbersome to execute, low in efficiency for generating positive roots, and vulnerable to explant contamination.

Betalains, the product of tyrosine-based substrate synthesis, are catalyzed by three enzymes: CYP76AD1, DODA, and GT. The Betalains synthesis system *Ruby* integrates the synthesis of these enzymes into a single open reading frame[20]. Given the ubiquitous presence of tyrosine in plant cells, *Ruby* has the theoretical potential for expression in any plant tissue, making it an ideal marker for positive root identification(He et al. 2020a; Ge et al. 2023).

In our study, we detail and illustrate the procedure for generating soybean hairy roots using cotyledons and soybean stems. When employing cotyledons as explants for hair root formation, a high success rate was observed: 90%-99% of the infected explants from four different cultivars successfully produced hairy roots, and among these, 30%-60% were transformed. In contrast, when utilizing stems as explants, up to 80% of the infected explants produced hairy roots in three different varieties. The transformation of soybean stems proved to be straightforward and does not necessitate a sterile environment. Remarkably, both systems collectively require only 22 days for the entire workflow and are compatible with a wide range of soybean genotypes. A key finding of this study is the effective use of the *Ruby* reporter gene for precise screening of positive roots. The establishment of an efficient in vitro hairy root system represents a rapid and effective platform for investigating gene function in soybeans. Transgenic hairy roots, generated through this system, are invaluable for various applications, including protein expression, subcellular localization studies, bimolecular fluorescent complementation (BiFC) analysis, and screening of target sgRNAs for CRISPR/Cas9 gene editing(Kong et al. 2023). Beyond these applications, the simplicity and efficiency of this soybean hairy root transformation method hold potential for broader implications in plant science(Ron et al. 2014; Cheng et al. 2021; Pereira et al. 2023). It can be adapted for exploring root biology in plant species beyond soybeans, offering a versatile tool for root-related research across a diverse range of plant genotypes.

## Results

### 1. Development of Three Distinct Reporter Vectors for Validating Soybean Transformation

The construction of several innovative selection marker constructs for soybean root transformation selection was explored due to the autofluorescence exhibited by soybean roots(Gijzen et al. 2008). Zsgreen and tdtomato were identified as more efficient in fluorescence compared to conventional GFP or RFP, offering potential advantages(Shaner et al. 2004; Nakamura et al. 2013). To leverage these advantages, we engineered two vectors specifically for expressing Zsgreen and tdtomato in soybean transformations. Our focus was also on utilizing the betalain visual reporter system in soybean root transformation. Adapting the Ruby reporter system from He et al., we developed a vector to co-express the betalain pathway genes *CYP76AD1*, *DODA*, and *Glucosyltransferase* effectively under a single promoter termed ‘Ruby 1+2+3’(He et al. 2020b). Additionally, we designed a vector, ‘Ruby1+2’, linking *CYP76AD1* and *DODA*, and a separate construct for *Glucosyltransferase* (’Ruby 3’). All reporter genes were driven by the 2x CaMV 35S promoter, with a separate transcription unit, replaceable through LIC cloning and assembled via the Golden Gate method (Figure 1A).

**Figure 1.**
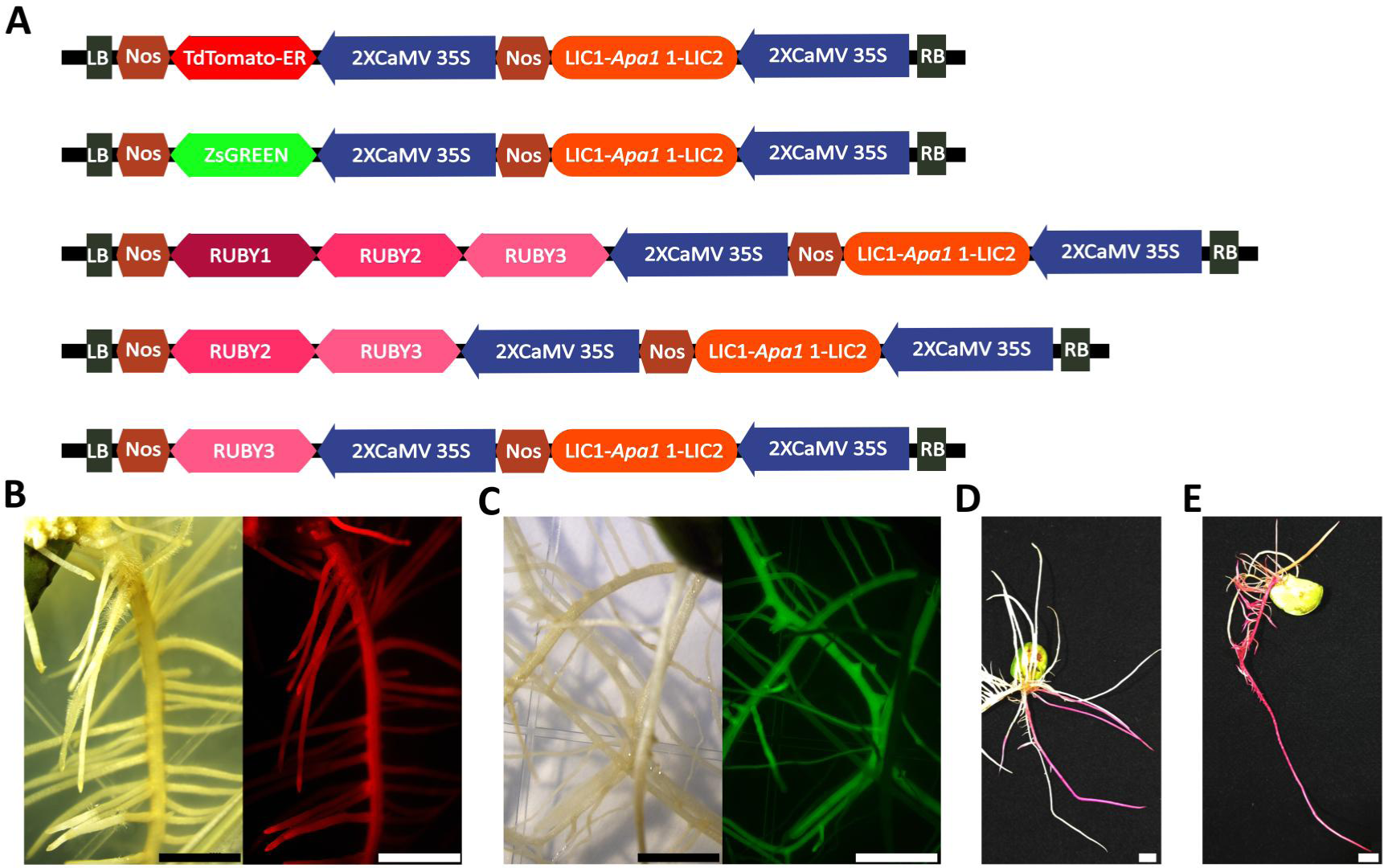
Assessment of Transformed Roots Using Specific Marker Genes. (A) Schematic representation of the TdTomato, ZsGreen, Ruby1+2+3, Ruby1+2, Ruby3. RUBY1, RUBY2, RUBY3 are different betalain biosynthetic genes. RUBY1 is *CYP76AD1*, RUBY2 is *DODA*, RUBY3 is *Glucosyltransferase*. (B) Roots transformed with TdTomato, shown under a stereomicroscope in bright field; scale bars = 4mm. The same TdTomato-transformed roots viewed under a stereomicroscope with a RFP filter set, exhibiting RFP fluorescence; scale bars = 4mm. (C) Roots transformed with ZsGreen, imaged under a stereomicroscope in bright field; scale bars = 4mm. The same ZsGreen-transformed roots visualized under a stereomicroscope with an GFP filter set, revealing GFP fluorescence; scale bars =4mm. (D) Display of hairy roots at 14 days post-inoculation (dpi) with *A. rhizogenes* Ar.Qual-*Ruby1+2+3*; scale bars =4mm. (E) Hairy roots shown at 14 dpi with *A. rhizogenes* Ar.Qual-*Ruby1+2* and Ar.Qual-*Ruby3*; scale bars =4mm.

Testing in soybean roots, mediated by *Agrobacterium rhizogenes*, revealed that Zsgreen and tdtomato were easily distinguishable from the natural root autofluorescence. Furthermore, the Ruby1+2+3 construct successfully produced enzymes for betalain synthesis. Co-delivery of Ruby1+2 and Ruby 3 by a mixture of two *Agrobacterium* strains resulted in the desired betalain red root phenotype. It was observed that both fluorescent proteins and betalain were uniformly expressed throughout the entire root system originating from the cotyledon (Figure 1B∼E).

### 2. SilwetL-77 and Hormone Mixtures in AAS could enhance transformation rate in hairy root transformation

Plant hormones are pivotal in root generation, and their inclusion during plant transformation is a widely recognized practice(Bahramnejad et al. 2019). Furthermore, incorporating detergents like Silwet L-77 has been shown to substantially enhance the efficiency of *Agrobacterium*-mediated plant transformation(Clough and Bent 1998). Based on these insights, we hypothesized that adding both plant hormones and Silwet L-77 during soybean root transformation could yield beneficial results. To test this, we refined the existing transformation procedure by introducing an additional step: immersing the wounded cotyledons in an *Agrobacterium* Auxiliary Solution (AAS) for 15 minutes during the soybean hairy root transformation process (Figure 2). This adjustment renders the soybean hairy root transformation procedure not only more comprehensive but also allows for the observation of induced transformed roots within approximately two weeks.

**Figure 2.**
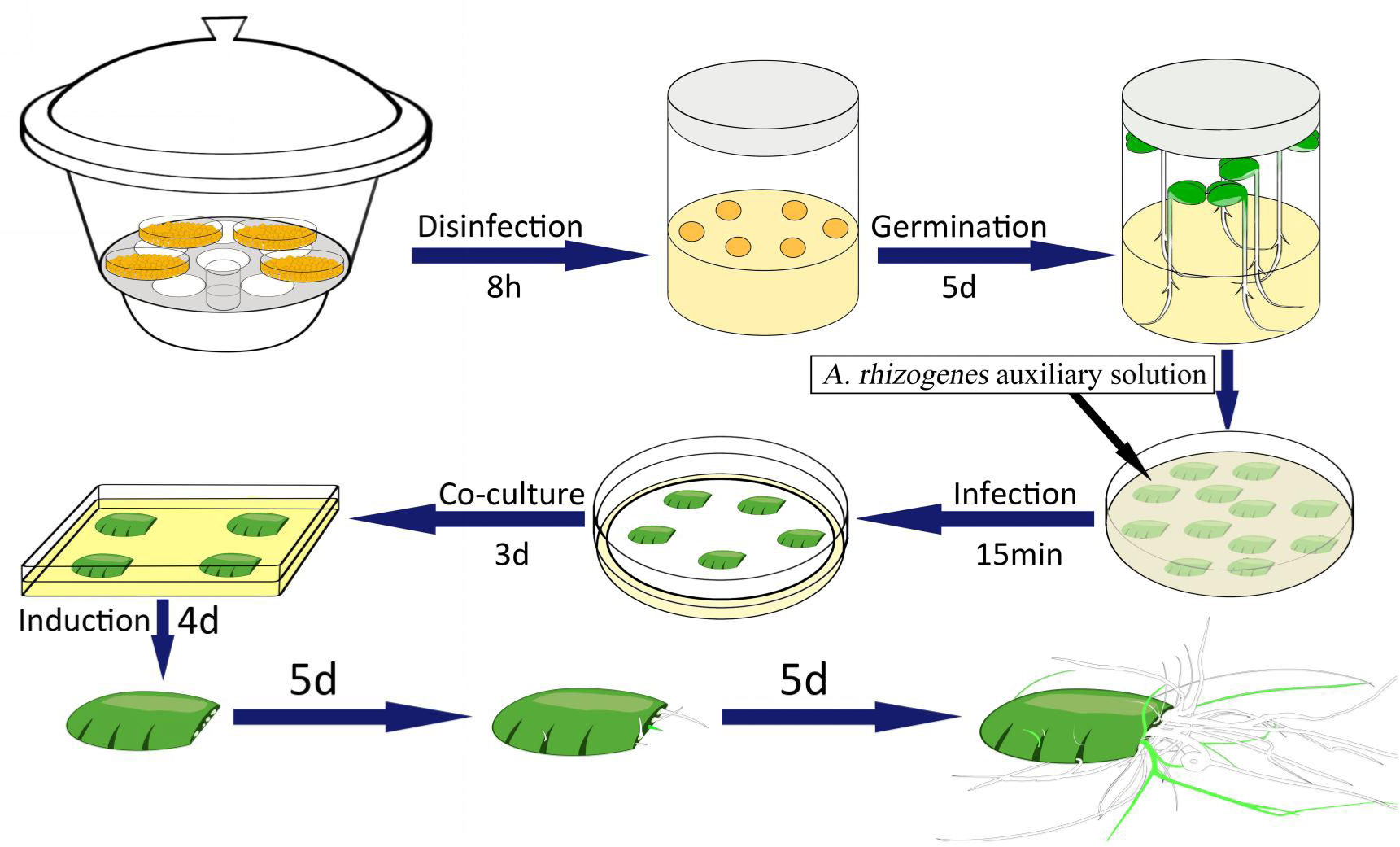
Schematic representation of the soybean hairy root transformation process using cotyledons as explants. The workflow begins with the disinfection of soybean seeds in a sealed desiccator for 8 hours. Subsequently, the seeds undergo a 5 day germination period. After germination, the cotyledons are immersed in AAS infection with *A. rhizogenes* for 15 minutes, followed by a 3-day co-culture period. The cotyledons are then transferred to an induction medium for 4 days duration, allowing induced hairy roots to develop over the subsequent 5 days. Finally, the roots are grown out for an additional 5 days to complete the transformation process, culminating in the production of transgenic soybean hairy roots.

Initially, we evaluated the effect of incorporating different hormone mixtures (HM) and a combination of hormone mixtures with Silwet L-77 (HM+SW) on the hairy root transformation efficiency in the soybean variety “Wandou28”. The results were quite insightful: adding only the plant hormone mixture to the *Agrobacterium* Auxiliary Solution (AAS) resulted in a marginal increase in total root transformation efficiency, approximately 1.9-fold (not statistically significant), based on the ratio of total positive roots to total roots generated from cotyledons. However, when the hormone mixture was combined with Silwet L-77, the transformation efficiency significantly increased to about 3.8 times compared to control (Figure 3A). We also assessed the cotyledon transformation rate, calculated as the ratio of cotyledons with positive roots to the total number of cotyledons used in a single experiment. The addition of the hormone mixture alone to the AAS led to a 2.5-fold increase in the cotyledon transformation rate. Furthermore, the inclusion of both HM and SW resulted in a 4.0-fold enhancement compared to the control group where no AAS was added (Figure 3B). From another perspective, we measured the total cotyledon root induction rate, defined as the ratio of cotyledons with induced roots to the total cotyledons. The introduction of HM alone showed no significant difference compared to the control, but the combined addition of HM and SW notably increased the cotyledon root induction rate by 1.4 fold (Figure 3C).

**Figure 3.**
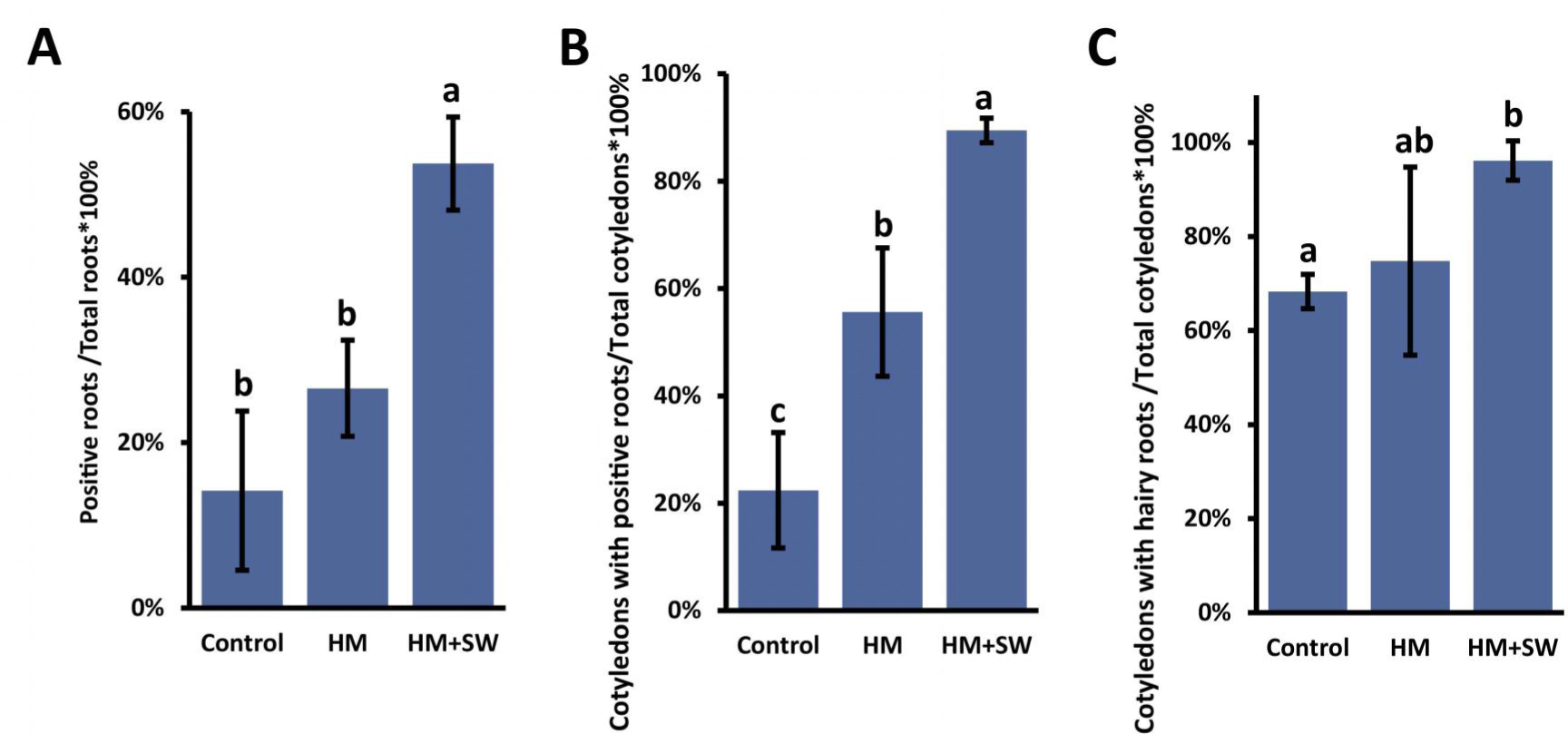
Comparative analysis of transformation efficiency influenced by AAS components. (**A**) quantifies the rate of positive roots compared to total roots, (**B**) assesses the proportion of cotyledons with positive roots, and (**C**) reflects the overall root induction rate as the ratio of cotyledons with hairy roots. Data points are the mean±SD from three biological replicates. Statistical significance is indicated by different letters, with distinct letters signifying significant differences and matching letters indicating no significant difference, as determined by Student’s t-test (P<0.05).

These results were consistent across various methods employed for assessing the formation of positive roots, indicating a reliable trend in efficiency improvement. The study clearly demonstrated that both the hormone mixture and the surfactant played significant roles in augmenting the conversion efficiency within the root development process. Notably, the most pronounced increase in transformation efficiency was observed when both the hormone mixture and the surfactant were used together, as illustrated in Figure.3. This synergy suggests that the combined use of these additives in the AAS could be a highly effective strategy for enhancing the hairy root transformation process in soybean research.

### 3. Enhanced Soybean Cotyledon Transformation with AAS across different varieties

In this detailed study, we evaluated the final positive transformation rates across several soybean varieties, including “Zhonghuang 39”, “Williams 82”, “Forrest”, and “Wandou28”. Our assessment focused on three key metrics: 1) total positive root transformation efficiency, calculated by the ratio of positive roots to total roots (Figure. 4A) cotyledon transformation efficiency, measured by the frequency of cotyledons developing positive roots and the total production of hairy roots from cotyledons (Figure. 4B) root induction rate, determined by dividing the number of cotyledons with induced roots by the total number of cotyledons (Figure. 4C).

**Figure 4.**
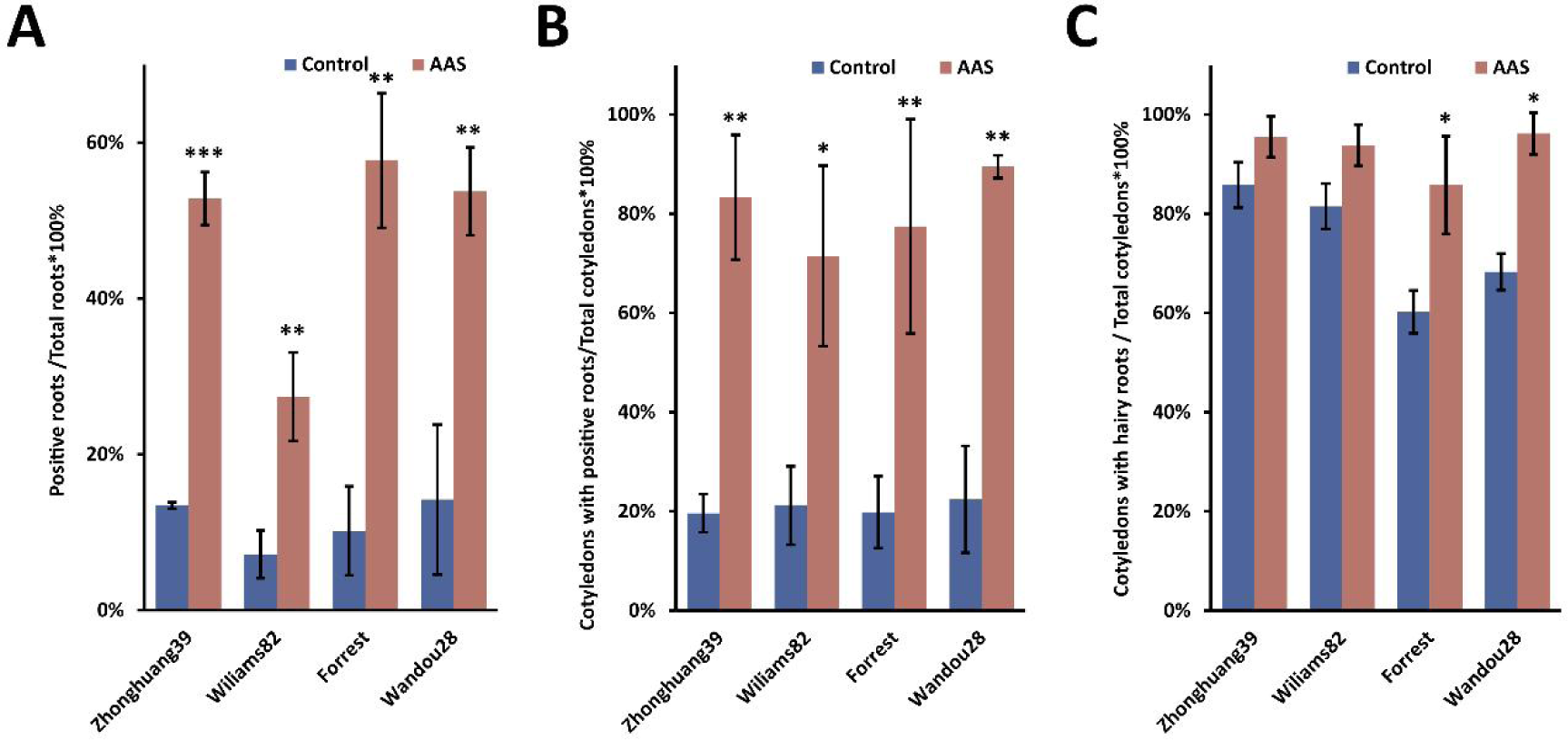
Genotypic impact on soybean transformation efficiency with AAS. Transformation efficiency in different soybean genotypes—’Zhonghuang39’ ‘Williams 82’ ‘Forrest’ and ‘Wandou28’—following the application of *A. rhizogenes* infestation AAS. **(A)** quantifies the rate of positive roots compared to total roots, **(B)** assesses the proportion of cotyledons with positive roots, and **(C)** reflects the overall root induction rate as the ratio of cotyledons with hairy roots. Data are presented as mean±SD from three biological experiments, with statistical significance denoted by asterisks as per Student’s t-test: *P<0.05, **P<0.01, ***P<0.001.

Our findings highlight that there was a notable increase in the rate of positive root conversions in each soybean variety, with efficiencies rising to 3-4 times higher than the baseline upon adding AAS (Figure.4A and 4B), however, incorporating AAS into the *A. rhizogenes* wound infestation process did not significantly impact the proportion of cotyledons producing hairy roots compared to the control group without AAS (Figure. 4C). Additionally, our research demonstrated a consistent boost in transformation efficiency across different soybean genotypes tested, indicating the broad applicability and efficiency of the hairy root transformation system. This uniformity emphasizes its potential as a dependable method for genetic research and biotechnological advancements in various soybean cultivars.

### 4. Significant Reduction in Transformation Rate with Larger Vector Sizes

In previous studies, vectors using markers like GFP (less than 1kb) or GUS (around 2kb) generally yielded high transformation rates. However, in our experiments, when the transformation vector was larger, such as containing an additional transcription unit or a marker significantly exceeding 2kb in size, we observed a lower transformation rate. To investigate this, we utilized a Ruby cassette with three genes totaling 4kb in length. We also tested a smaller variant, Ruby1+2, which is only 2.3 kb and results in a red root phenotype but is 1.7 kb shorter than the full Ruby1+2+3. Our findings indicate a noticeable difference in transformation efficiency: while markers like GFP and RFP showed root transformation efficiencies around 52%, the transformation rate for the Ruby1+2+3 reporter was significantly lower, at approximately 29% (Figure 5A). This trend was also evident in cotyledon transformations, where GFP and RFP achieved positive rates of about 89-96%, but the Ruby1+2+3 transformation rate was only 64%, notably lower (Figure 5B). Additionally, the rate of cotyledon root generation using the Ruby1+2+3 vector was slightly reduced, at 85% to 96%, compared to the GFP or RFP controls.

**Figure 5.**
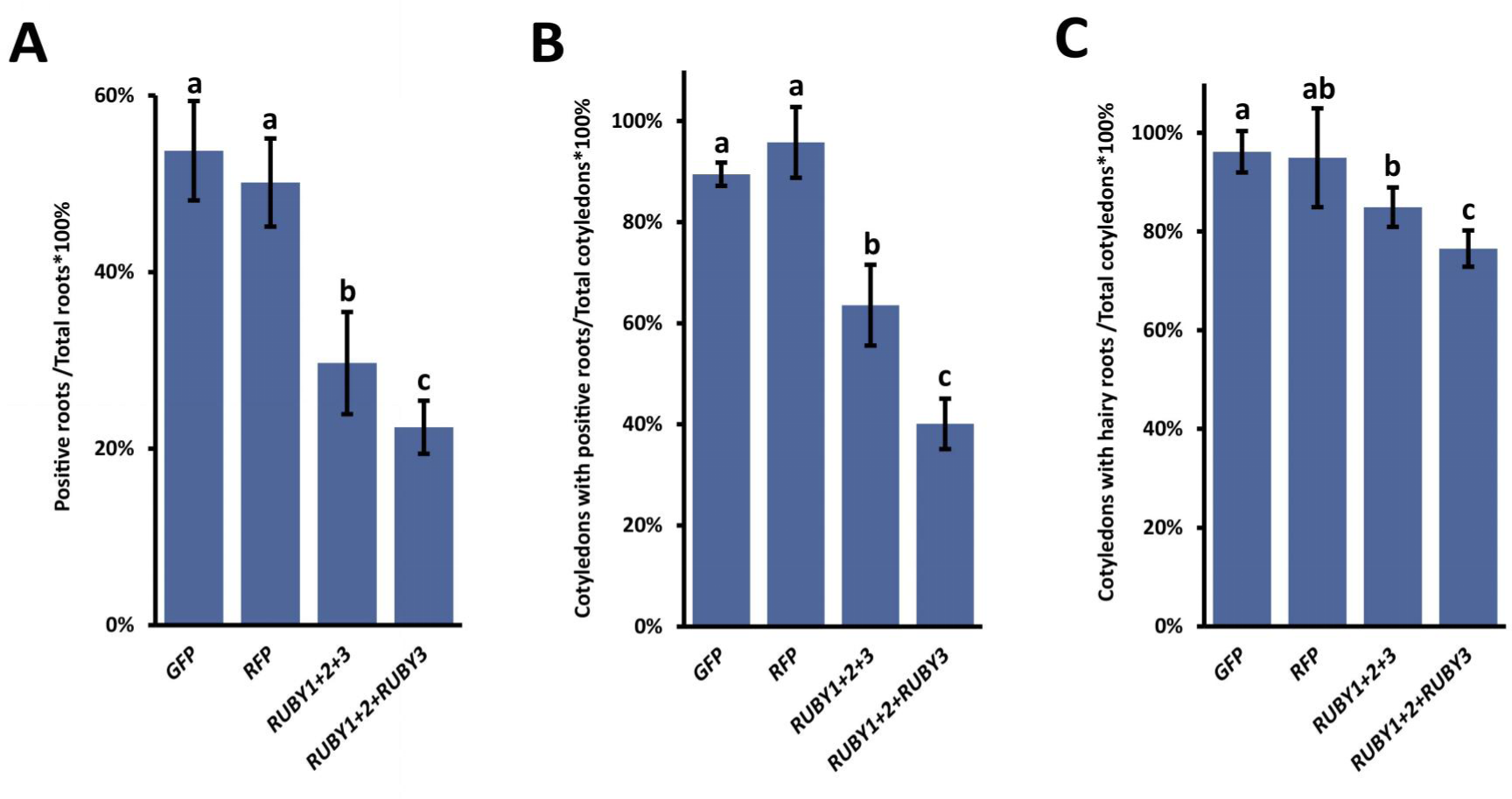
Comparative analysis of transformation efficiency influenced by reporter Genes. **(A)** quantifies the rate of positive roots compared to total roots, **(B)** assesses the proportion of cotyledons with positive roots, and **(C)** reflects the overall root induction rate as the ratio of cotyledons with hairy roots.Data points are the mean±SD from three biological replicates. Statistical significance is indicated by different letters, with distinct letters signifying significant differences and matching letters indicating no significant difference, as determined by Student’s t-test (P<0.05).

We also investigated the impact of AAS on the cotransformation rate of two separate vectors. For this purpose, we placed Ruby1+2 in one binary vector and Ruby3 in another. These two vectors, transformed in *A. rhizogenes* strains Ar.Qual, were mixed and transformed following the method described in Figure 2. The results indicated that successful cotransformation of both vectors, Ruby1+2 and Ruby 3, in a single cell was necessary to produce a functional marker, as evidenced by the red coloration of roots similar to that observed with the Ruby1+2+3 single vector (Fig.1D and 1E). Notably, the efficiency of cotransformation with the two vectors (Ruby1+2 and Ruby3) for both total root transformation and cotyledon transformation was slightly but significantly lower than that of the single vector transformation with Ruby1+2+3 (Figure 5A and 5B). Additionally, a lower percentage of cotyledons generated roots in the two-vector transformation with Ruby1+2 and Ruby3 compared to the single vector transformation with Ruby1+2+3.

In summary, our results clearly demonstrate that larger transformation vectors, such as the Ruby1+2+3 cassette, significantly reduce transformation efficiency in both root and cotyledon contexts compared to smaller markers like GFP and RFP. Furthermore, the addition of AAS was found to universally increase the transformation rate across all tested vector sizes for both total root and cotyledon positive cases.

### 5. Enhancing soybean hypocotyl transformation rate/ composite plants with AAS

Composite soybean plants are widely used in soybean research for their capability to generate transgenic roots through a simple, non-aseptic stem cut wound transformation process. We investigated if AAS could enhance the efficiency of hypocotyl transformation in composite soybean plants. Our procedure, adapted from a method outlined in (citation), involved immersing seedlings in an *A. rhizogenes* solution with AAS, followed by cutting the stem about 1 cm below the cotyledon. After around two weeks, we observed new transformed roots by removing the vermiculite from the wounded stem area (Figure 6).

**Figure 6.**
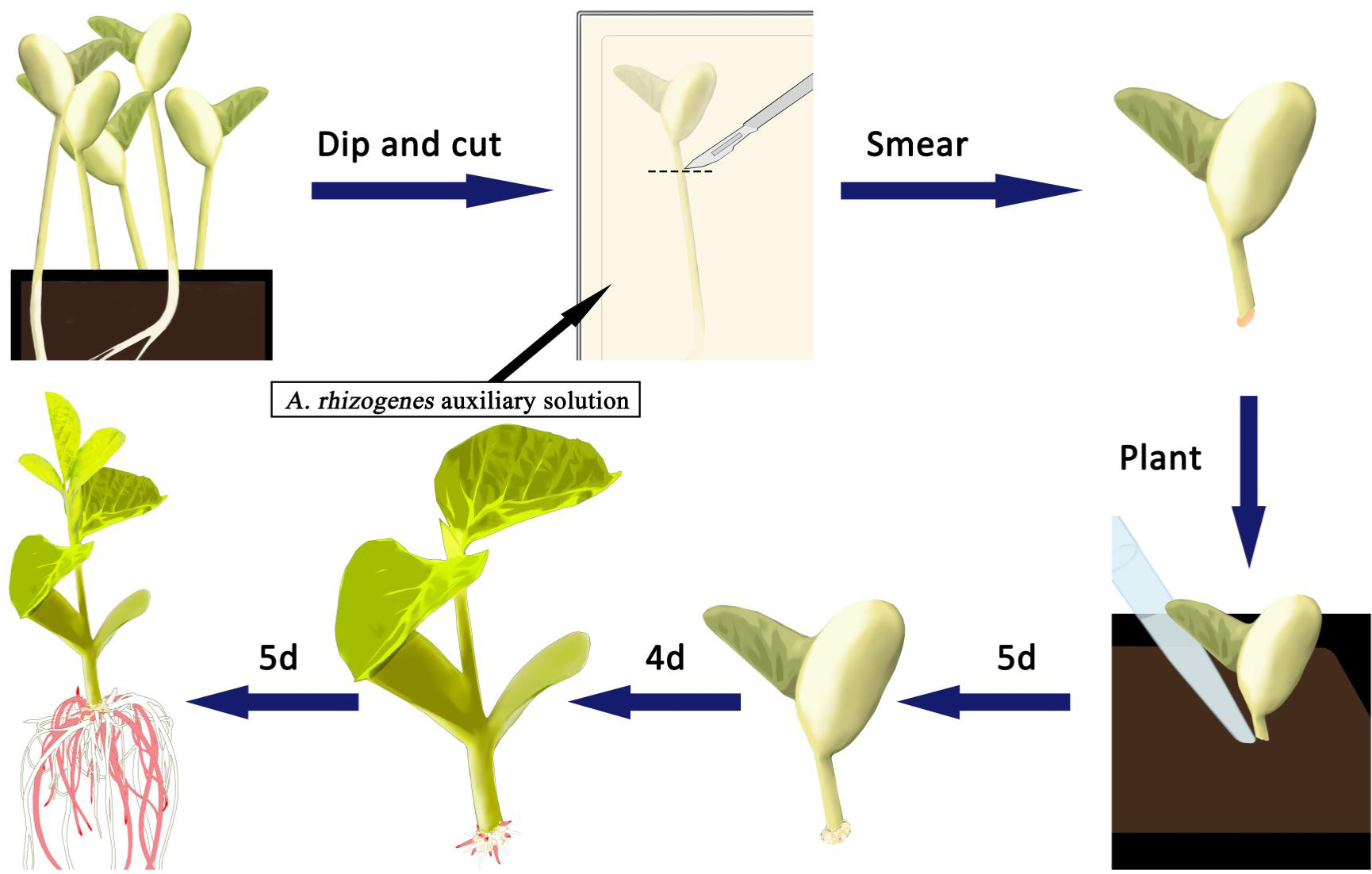
Schematic representation of soybean hairy root transformation process using stems. Soybean seedlings are cut near the roots after a six-day cultivation period. Seedling hypocotyls are submerged in the AAS and then cut 0.7cm-1cm from the cotyledons.The remaining hypocotyl after the cut is inoculated with *A. rhizogenes*. Seedlings are then planted in moist soil and irrigated with the AAS. Seedlings at 14 days post-inoculation (dpi), red roots are positive transformed roots.

For visual ease, we opted for *Ruby1+2* and *Ruby1+2+3* markers, both resulting in visibly red hairy roots under normal lighting conditions, making phenotype observation straightforward (supplemental figure 1). Notably, Ruby1+2 roots exhibited a lighter red (supplemental figure 1A) compared to the deeper pink of Ruby1+2+3 (supplemental figure 1B).

In this study, we evaluated the impact of AAS on hypocotyl transformation efficiency using three soybean varieties: “Williams 82”, “Forrest”, and “Dongnong 50”. We assessed transformation efficiency in two ways: positive root efficiency (number of positive roots per total roots generated) and positive explant efficiency (number of explants with positive roots per total explants). The results indicated that AAS generally enhanced both the positive root and explant efficiencies (figure 7 and table 1) across all tested varieties and both Ruby1+2 and Ruby1+2+3 vectors, aligning with observations in cotyledon hairy root transformation. Interestingly, while transformation efficiency and frequency Ruby1+2 were higher than with Ruby1+2+3, significant varietal differences in transformation rates were noted, particularly between “Forrest” and the other two varieties, “Williams 82” and “Dongnong 50”. Another finding of our study was the inability to achieve successful soybean hypocotyl transformation using *A. rhizogenes* Ar.Qual (data not shown). Despite extensive trials with the two reporter genes across multiple plant replicates, we did not observe any plants with red roots, indicating a lack of transformation success in this context. This outcome underscores the importance of choosing the right strains and conditions for effective soybean hypocotyl transformation.

**Figure 7.**
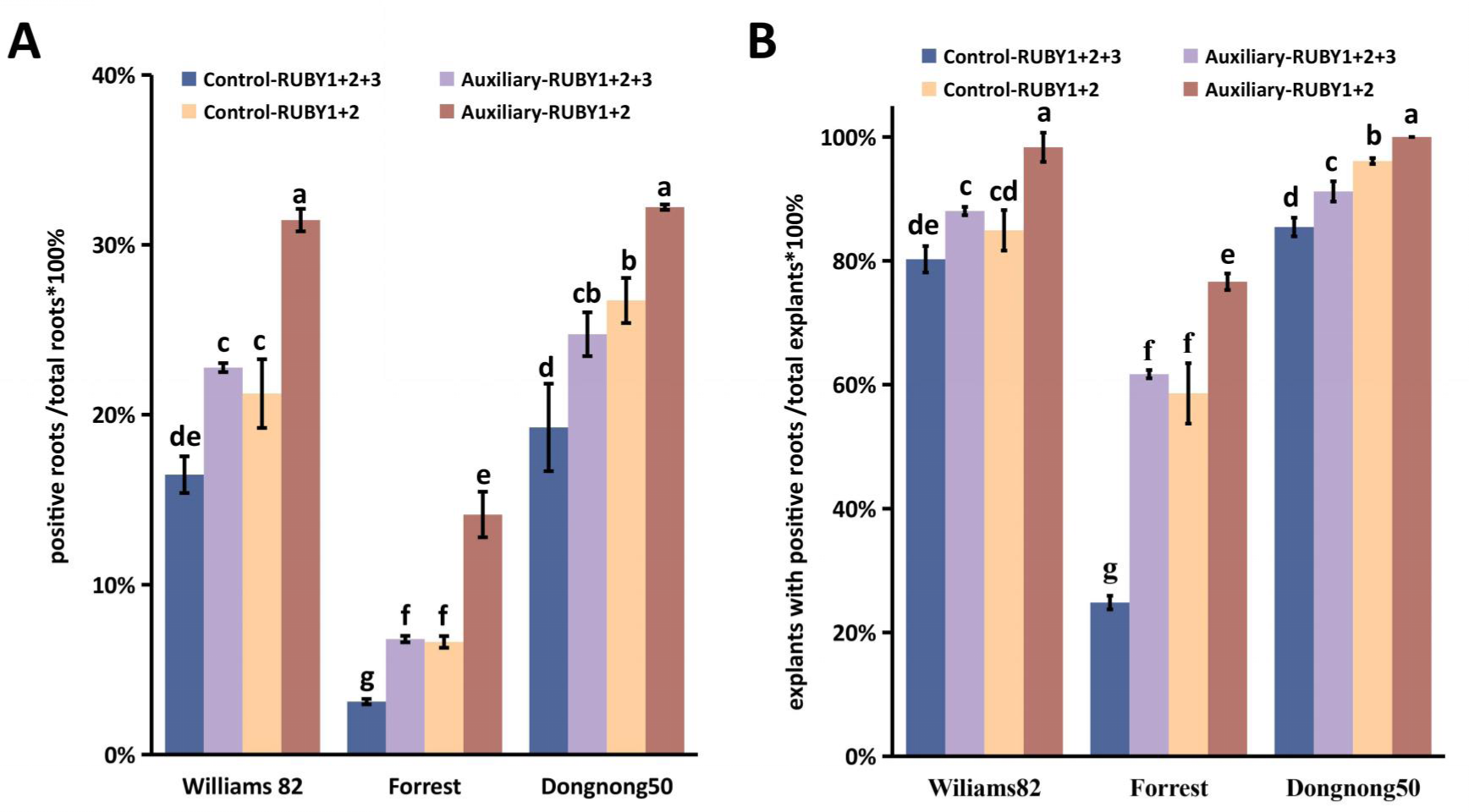
Soybean hypocotyl transformation efficacy across different genotypes and conditions. **(A)** Displays the transformation frequency, calculated as the percentage of positive roots relative to the total root count, across soybean genotypes ‘Williams 82’, ‘Forrest’, and ‘Dongnong50’ under different treatment conditions. **(B)** Illustrates the transformation efficiency, determined as the percentage of explants with positive roots out of all tested explants, for the same soybean genotypes and conditions. Data represent mean±SD from three biological replicates, with distinct letters indicating significant differences and identical letters indicating no significant difference as per the Student’s t-test (P<0.05).

**Figure 8.**
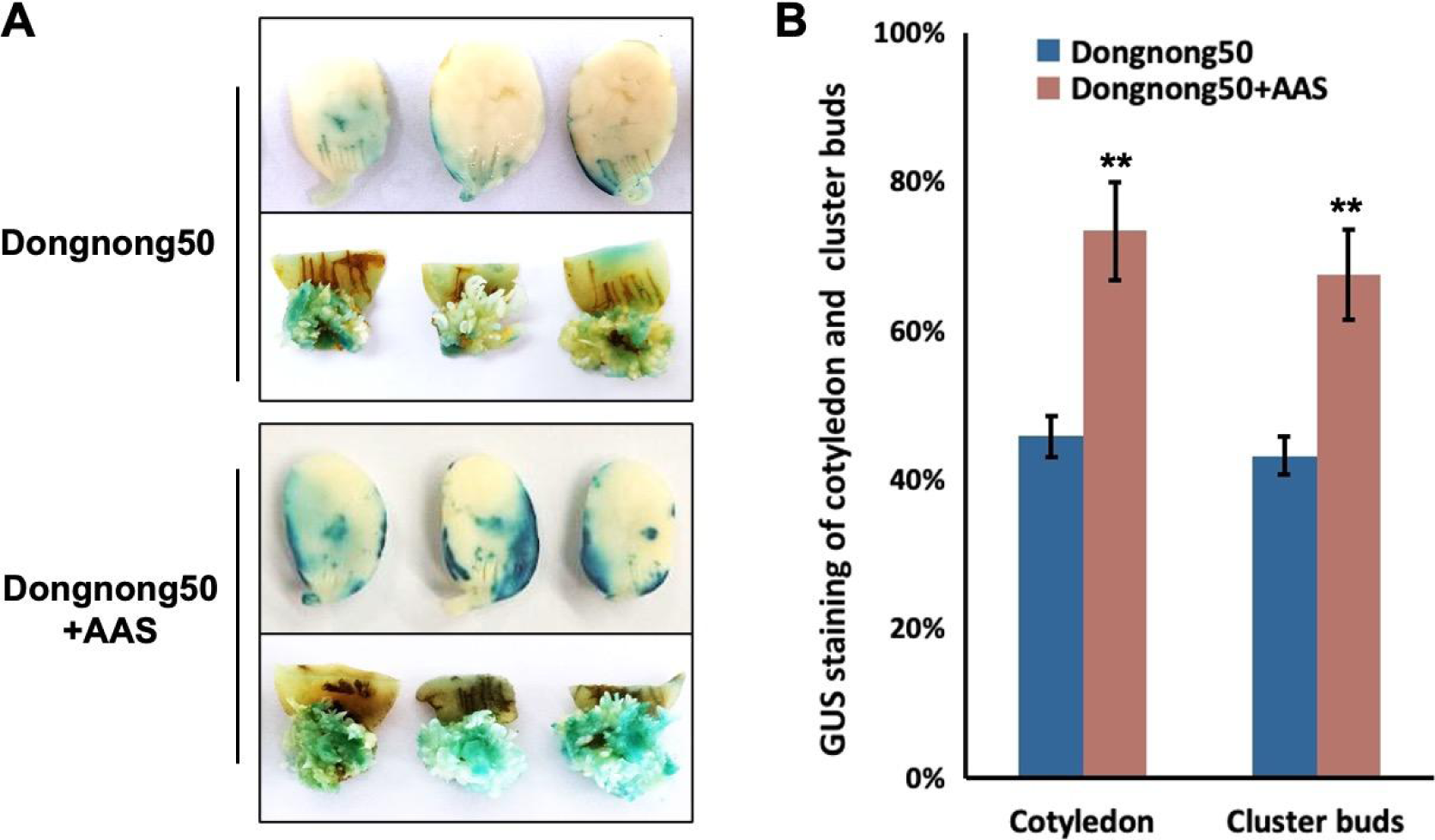
Enhancement in Soybean Cotyledonary Node Transformation Efficiency via Agrobacterium and the Addition of AAS. (A) A substantial increase in transformation efficiency is indicated by the GUS staining of cotyledons and clustered buds from Dongnong 50 following the addition of AAS. (B) Statistical analysis of GUS staining efficiency for cotyledons and clustered buds is shown. The data, expressed as mean±SD, are derived from three biological experiments, with statistical significance indicated by asterisks according to Student’s t-test: *P<0.05, **P<0.01.

**Table 1.** Outcomes of soybean hypocotyl transformation. The table summarizes the results of soybean hypocotyl transformation, detailing two key metrics. Transformation frequency is defined as the proportion of positive, transformed roots out of the total number of roots, expressed as a percentage. Transformation efficiency is calculated as the ratio of explants that developed positive, transformed roots to the total number of explants tested, also presented as a percentage.

| soybean<br>genotypes | Transformation<br>method | Numbers of<br>plants<br>infected | Numbers of the<br>explants with at least<br>one hairy root | Numbers of hairy<br>roots (per seedling) | Numbers of<br>positive roots<br>(per seedling) | Transformation<br>efficiency | Transformation<br>frequency |
| --- | --- | --- | --- | --- | --- | --- | --- |
| Williams 82 | 352-CK | 150 | 121 | 14.89±5.33 | 2.39±1.83 | 80.67% | 16.03% |
|  | 352-I | 150 | 132 | 14.21±5.25 | 3.27±2.20 | 88.00% | 23.02% |
|  | 353-CK | 150 | 128 | 13.92±4.93 | 3.32±2.24 | 85.33% | 23.82% |
|  | 353-I | 150 | 149 | 14.35±3.96 | 4.35±2.05 | 99.33% | 30.31% |
| Forrest | 352-CK | 150 | 37 | 14.00±4.52 | 0.45±0.87 | 24.67% | 3.19% |
|  | 352-I | 150 | 93 | 14.13±3.49 | 0.96±0.92 | 62.00% | 6.78% |
|  | 353-CK | 150 | 89 | 14.48±4.67 | 0.96±0.97 | 59.33% | 6.61% |
|  | 353-I | 150 | 115 | 13.93±4.33 | 2.01±1.95 | 76.67% | 14.44% |
| Dongnong 50 | 352-CK | 150 | 128 | 13.46±4.43 | 2.51±2.02 | 85.33% | 18.65% |
|  | 352-I | 150 | 137 | 13.81±3.48 | 3.39±2.47 | 91.33% | 24.55% |
|  | 353-CK | 150 | 144 | 13.56±4.63 | 3.45±2.16 | 96.00% | 25.41% |
|  | 353-I | 150 | 150 | 13.07±3.35 | 3.96±2.02 | 100.00% | 30.32% |

In conclusion, our research demonstrates that the addition of AAS significantly enhances hypocotyl transformation efficiencies in composite soybean plants, with marked improvements observed across various soybean varieties and vector types. This finding, coupled with the distinctive transformation rates among different varieties, underscores the potential of AAS in optimizing genetic modification techniques in soybean research.

### 6. Enhancing Efficiency of Agrobacterium-Mediated Transformation of Soybean Cotyledonary Node method through the Addition AAS

To assess the efficacy of AAS in improving Agrobacterium-mediated transformation efficiency in soybean cotyledonary nodes, we conducted experiments using the pCAMBIA3301-GUS vector, which includes the GUS reporter gene regulated by a constitutive promoter. Our findings reveal that AAS significantly enhances the transformation efficiency. Specifically, with the addition of AAS, the GUS staining percentage in Dongnong 50 cotyledons soared from 45.8% to 73.4%, demonstrating a notable increase in efficiency.

Moreover, the percentage of GUS staining in adventitious shoots rose from 43.2% to 67.5% with AAS treatment, further validating the positive impact of AAS on transformation success. These results are in line with previous research that has shown surfactants to significantly enhance Agrobacterium-mediated transformation in various plant species.

## Discussion

The results of our study provide compelling evidence of the pivotal role AAS play in enhancing the transformation efficiency of soybean plants, particularly in the context of hairy root and hypocotyl transformation processes. The innovative use of reporter vectors such as ZsGreen, TdTomato, and the Ruby series in soybean transformation validation offers a vivid and effective means to monitor and assess the success of genetic modifications.

The marked improvement in transformation rates observed with the inclusion of Silwet L-77 and hormone mixtures in AAS underscores the significance of optimizing the transformation medium’s composition. The synergy between these components in enhancing the hairy root transformation process is particularly notable, as it represents a promising avenue for maximizing transformation efficiency in soybean research.

Furthermore, our findings reveal the critical influence of vector size on transformation efficiency. The significant reduction in transformation rates with larger vectors underscores the need for careful consideration of vector design in the development of transformation strategies. This observation is crucial, especially in the context of complex genetic modifications where multiple genes are introduced.

The application of AAS in the generation of composite soybean plants has proven to be a game-changer, significantly boosting hypocotyl transformation rates across various soybean varieties. The positive impact of AAS on both the root and explant transformation efficiencies highlights its universal applicability and potential to revolutionize soybean genetic research.

However, our study also indicates the challenges associated with soybean hypocotyl transformation using certain strains of *A. rhizogenes*, as demonstrated by the unsuccessful attempts with the Ar.Qual strain. This finding emphasizes the necessity of strain selection and optimization of transformation conditions to achieve successful outcomes.

In light of these results, it is evident that the integration of AAS into soybean transformation protocols presents a significant advancement in the field. The enhanced transformation efficiencies, coupled with the ability to monitor and validate genetic modifications effectively, pave the way for more sophisticated and reliable genetic studies. The insights gained from this research not only enrich our understanding of soybean transformation dynamics but also hold profound implications for the broader field of plant biotechnology.

## Materials and Methods

### Plant materials and growth conditions

In this research, five soybean varieties—Williams 82, Zhonghuang 39, Forrest, Wandou 28, and Dongnong 50—were sourced from the Chinese Academy of Agricultural Sciences in Beijing and Zhejiang University. The soybean seeds were sprouted and cultivated in soil within a greenhouse, subject to long-day lighting conditions (16 hours of light followed by 8 hours of darkness), at a temperature of 25℃ and a humidity level of 60%.

### *Agrobacterium* Strains and Vector Constructions

The strains Ar.Qual and K599 of Agrobacterium rhizogenes, along with Agrobacterium tumefaciens strain EHA105, sourced from Weidi Bio under the catalog numbers AC1060, AC1080, and AC1010 respectively, were employed to promote the development of transgenic roots and cotyledonary nodes in soybeans. The *ZsGreen/GFP* and *TdTomato/RFP* genes were individually integrated into the Golden Gate binary vector pAGM4673, each controlled by the double cauliflower mosaic virus (CaMV) 35S promoter and terminated with a Nos terminator. The genes *Ruby 1+2* and *Ruby 3*, initially derived from the original Ruby vector pDR5:RUBY as a gift from Yubing He, were inserted into vector pAGM4673. This insertion was also under the control of the double CaMV 35S promoter and terminated with a Nos terminator, utilizing Golden Gate assembly techniques.

### Cotyledon Transformation Method

The transformation techniques were modified from the methods described by Han et al.(Han et al. 2023), with the incorporation an additional enhancement using AAS while maintaining the same medium composition otherwise. The soybean cotyledon transformation commenced with the sterilization of soybean seeds using chlorine gas. The seeds were germinated for 5 days on a specific medium, exposed to long-day conditions (16 hours of light and 8 hours of darkness) at a temperature of 25℃.. A technical *A. rhizogenes* solution was prepared by introducing a plasmid carrying the target gene in the *A. rhizogenes* Ar.Qual strain, followed by cultivation in LB medium. This bacterial pellet was subsequently suspended in an AAS specifically formulated with B5 medium, containing with 30g/L sucrose, 3.9 g/L MES sodium salt, 100 µl/L Silwet L-77, 40 mg/L Acetosyringone, 1.67 mg/L 6-Benzylaminopurine (6-BA), and 0.025 mg/L Gibberellin A3. During the infection process, the seed coats and true leaves of the germ-free seedlings were excised, the cotyledons were incised and soaked in the AAS mixture for 15 minutes. The treated cotyledons were then placed on a co-culture medium and kept in darkness for 3 days. Root initiation was stimulated by transferring the cotyledons onto a rooting medium, with hairy root development typically visible after 14 days. The growth of hairy roots was subsequently verified using a fluorescence microscope, identifying positive roots through GFP or RFP filters.

### Soybean hypocotyl transformation method

In the soybean hypocotyl transformation method, germination involves spreading seeds on moistened coarse vermiculite in a seedling tray, covered and cultivated for 6 days under a light-dark cycle(16 h light/8 h dark) at 25°C. For the AAS, the target gene plasmid is transformed into *A. rhizogenes* K599, and single colonies were cultured in LB medium. The *A. rhizogenes* pellet was subsequently reconstituted in the AAS, identical to that outlined in the section describing the Cotyledon Transformation Method. Infection is done by cutting near the roots(cut 0.7cm-1cm from the cotyledons) and soaking the stems in this solution. The remaining hypocotyl after the cut is inoculated with *A. rhizogenes.* The inoculated seedlings were planted in moist soil and covered to maintain humidity, fostering hairy root development over a 14 day period. Positive roots, marked by the presence of *Ruby* screening genes, are identified as red roots after washing off the soil.

### Transformation of Soybean Cotyledonary Nodes Mediated by *Agrobacterium tumefaciens*

In the transformation study, the pCAMBIA3301-GUS vector with a GUS reporter gene was utilized. We infected soybean cotyledonary nodes of the Dongnong 50 variety with Agrobacterium tumefaciens strain EHA105, containing this vector. The infection solution was augmented with AAS to examine its effect on infection efficiency. Nodes were immersed in an Agrobacterium suspension, with and without AAS, and agitated at 28°C for 30 minutes.

This process involved 200 explants per treatment in three replicates. After 3 days of co-cultivation and 14 days of bud induction, nodes were stained, vacuum infiltrated at 70 kPa for 30 minutes, and incubated overnight at 37°C. Decolorization followed using an alcohol gradient (95%, 70%, 50%), involving 50 nodes per treatment in three replicates. GUS staining was then applied to assess transgene expression and transformation efficiency.

## Acknowledgements

We thank Dr. Yubing He for providing Ruby original plasmids. We thank Dr. Ray Collier for sharing Golden Gate cloning parts and advice. We thank Dr. Shiming Liu and Dr. Lingan Kong for sharing soybean varieties and advice. This work was funded by the National Key Research and Development Program of China (2023YFD1401000), the National Natural Science Foundation of China (32272478 and 32102146), Natural Science Foundation of Zhejiang Province (LTGN23C130003).

## Author contributions

L.C. designed the research, performed research, analyzed data and wrote the paper. L.W. performed research and analyzed data. S.H. and Y.L. designed the research, analyzed data and wrote the paper.

---

**Supplemental Figure 1.**
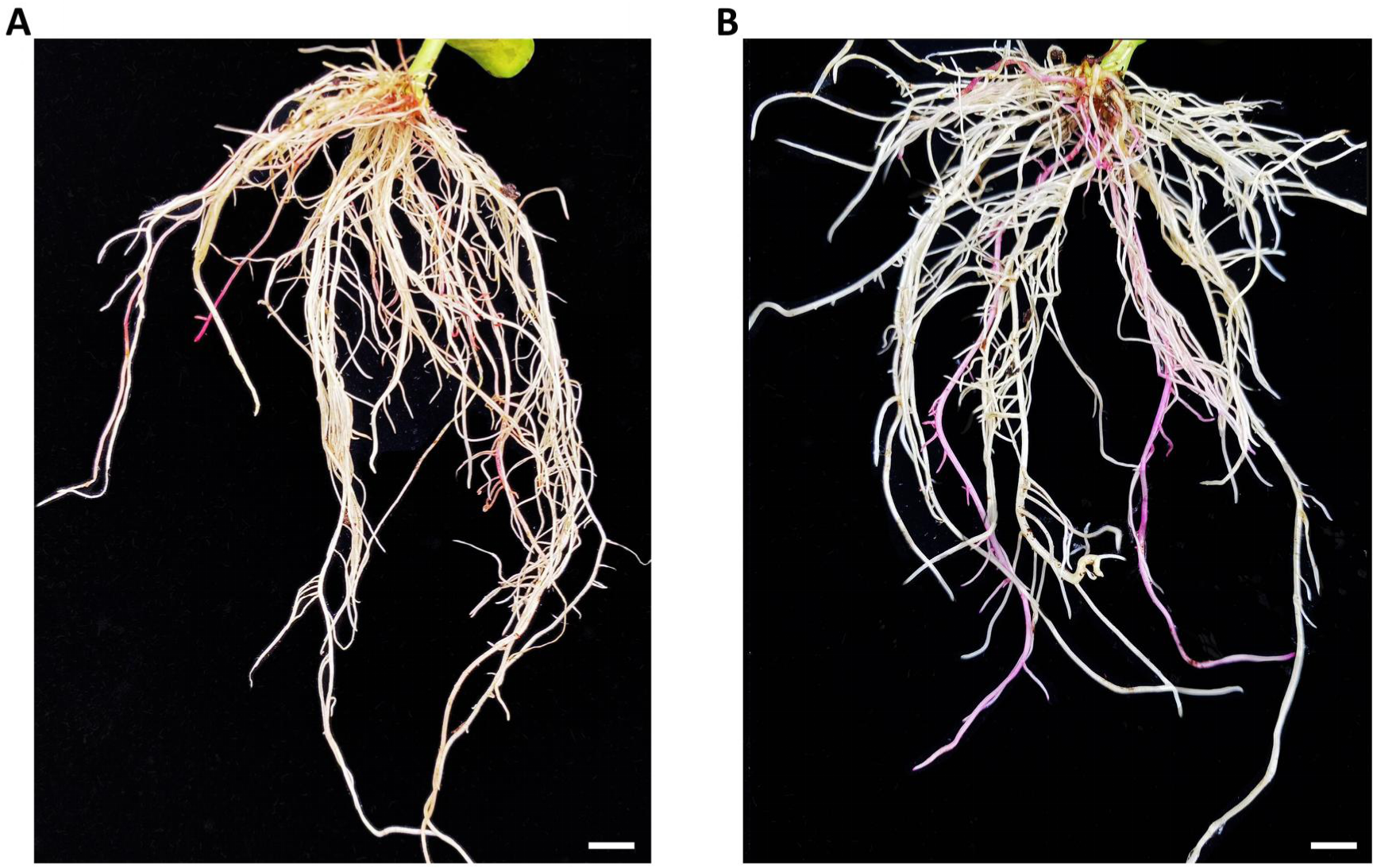
Genotypic impact on soybean transformation efficiency with AAS. **(A)** At 14 days post-inoculation (dpi), hairy roots are induced transformed with 35s::Ruby1+2+3 plasmids in *A. rhizogenes* K599. red roots are positive transformed roots; scale bars = 4mm. **(B)** Hairy roots are similarly induced at 14 dpi transformed with 35s:Ruby1+2 plasmids in *A. rhizogenes* K599. pink roots are positive transformed roots; scale bars = 4mm.

